# Characterization of adenine phosphoribosyltransferase (APRT) activity in *Trypanosoma brucei brucei*: Only one of the two isoforms is kinetically active

**DOI:** 10.1101/2021.10.20.465077

**Authors:** Kayla Glockzin, Thomas D Meek, Ardala Katzfuss

## Abstract

Human African Trypanosomiasis (HAT), also known as sleeping sickness, is a Neglected Tropical Disease endemic to 36 African countries, with approximately 70 million people currently at risk for infection. Current therapeutics are suboptimal due to toxicity, adverse side effects, and emerging resistance. Thus, both effective and affordable treatments are urgently needed. The causative agent of HAT is the protozoan *Trypanosoma brucei* ssp. Annotation of the *T. brucei* genome confirms previous observations that *T. brucei* is a purine auxotroph. Incapable of *de novo* purine synthesis, these protozoan parasites rely on purine phosphoribosyltransferases to salvage purines from their hosts for the synthesis of purine monophosphates. Complete and accurate genome annotations in combination with the identification and characterization of the catalytic activity of purine salvage enzymes enables the development of target-specific therapies in addition to providing a deeper understanding of purine metabolism in *T. brucei*. In trypanosomes, purine phosphoribosyltransferases represent promising drug targets due to their essential and central role in purine salvage. Enzymes involved in adenine and adenosine salvage, such as adenine phosphoribosyltransferases (APRTs, EC 2.4.2.7), are of particular interest for their potential role in the activation of adenine and adenosine-based pro-drugs. Analysis of the *T. brucei* genome shows two putative *aprt* genes: APRT1 (Tb927.7.1780) and APRT2 (Tb927.7.1790). Here we report studies of the catalytic activity of each putative APRT, revealing that of the two *T. brucei* putative APRTs, only APRT1 is kinetically active, thereby signifying a genomic misannotation of Tb927.7.1790 (putative APRT2). Reliable genome annotation is necessary to establish potential drug targets and identify enzymes involved in adenine and adenosine-based prodrug activation.

**Author Summary:** Neglected Tropical Diseases, are defined by the World Health Organization as a diverse group of 20 different diseases that disproportionally affect the world’s poorest populations, including Human African Trypanosomiasis (HAT). HAT is endemic to 36 African countries and approximately 70 million people worldwide are currently at risk for infection. Current therapeutics are suboptimal due to toxicity, adverse side effects, and emerging resistance. The causative agent of HAT is *Trypanosoma brucei* ssp. Unlike humans, these protozoan parasites rely on purine phosphoribosyltransferases to salvage purine bases from their hosts for the synthesis of DNA and RNA. Here we report on the characterization of adenine phosphoribosyltransferase (APRT) activity in *Trypanosoma brucei brucei*. Our studies reveal that of the two putative APRTs, only APRT1 is kinetically active under physiological conditions. Accurate genome annotations in combination with the characterization of purine salvage enzymes allows the development of target-specific therapies. Enzymes involved in adenine and adenosine salvage, such as APRTs, are of particular interest for their potential role in the activation of adenine and adenosine-based pro-drugs.

## Introduction

*Trypanosoma brucei* ssp. is the etiological agent of Human African Trypanosomiasis (HAT), also known as sleeping sickness. *Trypanosoma brucei gambiense* and *Trypanosoma brucei rhodesiense* cause, respectively, chronic and acute forms of this disease in humans [1, 2], for which the chronic infection is more prevalent (> 90% of reported cases) [3]. HAT presents initial symptoms of headache, fever, and fatigue; and is mainly transmitted horizontally by the vector tsetse flies of the genus *Glossina* [4], but also vertically from mother to offspring [5], with evidence of sexual transmission [6]. Once in the central nervous system (CNS), the parasites alter sleeping habits, cause mood swings, impairment of reasoning and mental processing, followed by coma and death if untreated [7-9]. Progression of HAT leads to demyelination in the CNS, making it impossible to cure even after clearance of parasites [1]. HAT is endemic to 36 African countries; threatening an estimated 70 million people [3, 10].

No vaccine or prophylactic therapy is available [8], and the current standard of treatment is poor. Only four drugs are currently used to treat HAT. Pentamidine and Suramin are used in early stages of the infection. Melarsoprol is the first-line treatment for late-stage disease; and Eflornithine is effective only against *T. brucei gambiense* infections, but the dosage necessary limits its use. Treatment failure due to patient non-compliance ranges around 30%, with high rates of death related to toxicity of Melarsoprol (up to 12%) [1, 3]. The World Health Organization has emphasized the need for intensive efforts to treat HAT, including the development of novel therapies [11]; nonetheless, the last drug introduced for treatment was Eflornithine in 1990 [3].

Difficulties in the development of novel drugs for the treatment of Neglected Tropical Diseases include many social, economic, and political factors. In the case of HAT, these complications are exacerbated by the intrinsic complexities of the *T. brucei* ssp. convoluted life cycle. *T. brucei* ssp. is a single-cell obligate parasite with human and insect hosts. To complete its life cycle, the parasite must adapt to the widely different environments it faces in both hosts, and it accomplishes this task by modulating its gene expression, metabolic state, morphology, and reproduction strategies [4, 12-14]. Inside the tsetse fly, procyclic forms of trypanosomes adapt to a range of microenvironments including the proventriculus, salivary glands, anterior midgut and posterior midgut [15-17]. In the human host, the bloodstream forms of *T. brucei* ssp. infect blood and interstitial fluids, including cerebrospinal fluid and lymph [18].

The genome sequencing of *T. brucei* was completed in 2005 for the *T. brucei brucei* strain TREU927. *T. brucei brucei* causes disease (Nagana) in cattle and other vertebrates [19], however it is incapable of infecting humans and is considered the model organism for the infectious subspecies [20-22]. The data derived from the genomic annotation, allied to several proteomics studies that identified the differentially expressed genes along *T. brucei*’s life cycle [23-26] allows the use of target-specific strategies of drug development as a promising way to identify and develop novel therapies [27]. Genomic annotation of *T. b. brucei* corroborate previous observations that, as other parasitic protozoa, *T. brucei* ssp. is a purine auxotroph [27, 28], being incapable of synthesizing purine nucleotides *de novo*, and relying on uptake from the hosts to afford its metabolic needs via the purine salvage pathway. Trypanosomes incorporate purine nucleobases and purine analogs by a highly selective nucleobase/proton symporter system with differential expression during procyclic and bloodstream stages [29-32]. Once inside the cell, the purine nucleobases and nucleosides are further converted into nucleotides by enzymes of the salvage pathway, including three purine phosphoribosyltransferases which have some overlapping substrate specificity. Many of the enzymes of purine salvage present stage-specific expression profiles [33]; and previous cellular studies indicate that any single purine can be sufficient to support *T. brucei* growth *in vitro* [34]. Proper genome annotation, and the identification and characterization of the catalytic activity of purine salvage enzymes are key steps not only for understanding *T. brucei* metabolic pathways, but to aid the development of target-specific novel therapies. The enzymes involved on adenine and adenosine salvage are of particular interest for their potential role in the activation of adenine and adenosine-based antimetabolite pro-drugs [35-38].

In this work, we describe the kinetic characterization of *T. brucei brucei* adenine phosphoribosyltransferase (APRT) activity. APRTs (EC 2.4.2.7) are enzymes of the purine salvage pathway, catalyzing the formation of adenine monophosphate (AMP) from 5-phospho-α-D-ribose-1-diphosphate (PRPP) and adenine (Fig 1). A previous phenotypic study of APRT activity in *T. brucei* was conducted with an APRT1/APRT2 double knockout [35]. Other reports on APRT were performed using total cell extract activity [39], detection of mRNA [24], co-immunoprecipitation [40], or proteomic analyses [26]. The APRTs have been evaluated as potential drug targets, however previous work has been unsuccessful in evaluating APRT1 and APRT2 independently [41]. In this study, we focus on the catalytic activity of each putative APRT individually. Our results reveal that of the two *T. brucei brucei* genes annotated as putative APRTs, only one, APRT1, is biologically relevant for catalyzing the conversion of adenine into AMP.

**Fig 1.**
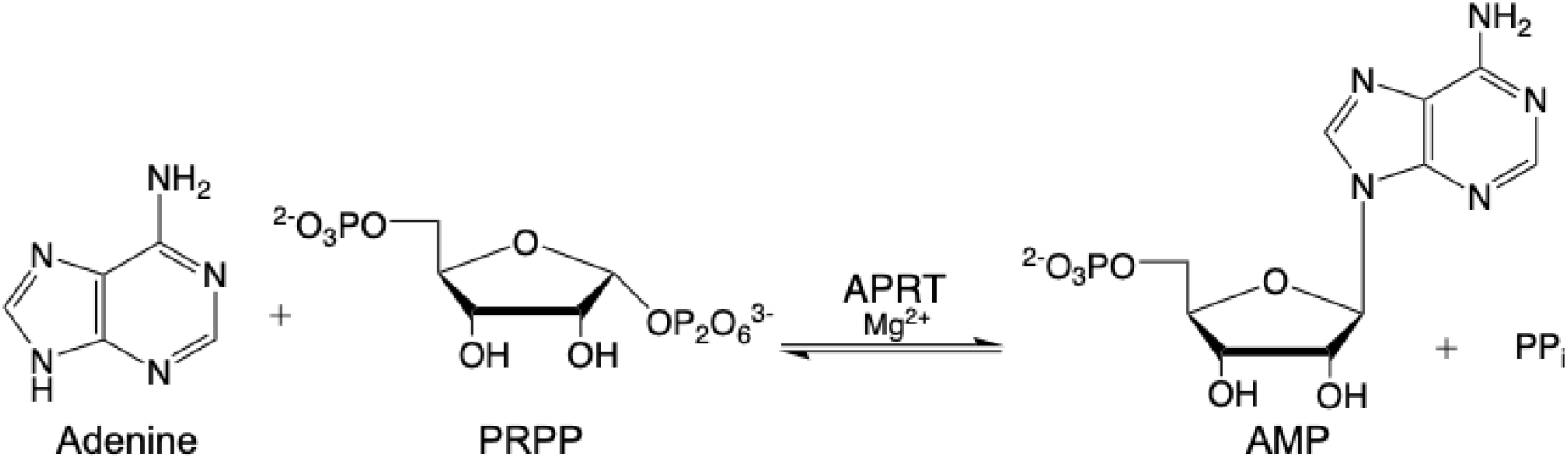
Reaction catalyzed by adenine phosphoribosyltransferase (APRT, EC 2.4.2.7).

## Methods

### Protein cloning, expression, and purification

APRT1 (Tb927.7.1780, UNIPROT access code Q57V32) and APRT2 (Tb927.7.1790, UNIPROT access code Q57V31) were codon optimized for expression in *Pichia pastoris*. Overhangs were added to the 5’ and 3’ ends of each ORF for assembly into the pPICZ vectors (S1 Appendix). Each gene was initially cloned into pET-28a(+) vectors, using *Nde*I and *Hind*III restriction sites (Genscript, USA). The plasmids pET-28a(+)::*aprt1-Ntag*, pET-28a(+)::*aprt1-Ctag*, pET-28a(+)::*aprt2-Ntag* and pET-28a(+)::*aprt2-Ctag* were linearized by PCR (S2 Appendix) and cloned into the pPICZ vectors (pPICZ-Ntag and pPICZ-Ctag) using a HiFi DNA assembly kit (NEB). Assembled products were subsequently transformed into NEB 5-alpha competent *Escherichia coli* cells using heat-shock and incubated overnight at 37°C on LB agar plates containing kanamycin 50 μg mL^-1^. Plasmids were linearized by PCR prior to transformation into *P. pastoris* strain smPP, using primers forward (5’ to 3’) GCTGTCTTGGAACCTAATATG, and reverse (5’ to 3’) TGTCAGTTTTGGGCCATTTG. The pPICZ-Ctag vector contains a cleavage site for the Tobacco Etch Virus (TEV) protease upstream of a C-terminal mCherry tag followed by a Strep-II tag and 6×His-tag. The pPICZ-Ntag vector contains an N-terminal TEV protease cleavage site downstream from a 6×His-tag followed by a mCherry tag (S1 Fig and S2 Fig). The clones with the highest expression levels of APRT1 and APRT2 were selected based on mCherry expression levels (S3 Fig). One single colony was used to inoculate 25 mL of YPD media in the presence of zeocin 50 μg mL^-1^. Pre-starting cultures were incubated overnight at 30°C in an orbital shaker set at 200 RPM. A 10 mL sample of pre-starting culture was used to inoculate 600 mL of BMGY media in the presence of zeocin 50 μg mL^-1^. Starter cultures were grown at 30°C and 200 RPM for 24 hours and harvested by centrifugation 2,000 × *g*, 5 min, at room temperature (RT). Pellets were resuspended with 40 mL of fresh BMMY media. 10 mL samples of the resuspended pellet were used to inoculate 600 mL of BMMY media in the presence of zeocin 50 μg mL^-1^. Cultures were grown at 30°C and set at 200 RPM for 24 hours. Protein expression was induced by addition of methanol at a final concentration of 0.5% (v/v). Cells were further grown under these conditions for 24 hours. Cells were harvested by centrifugation at 2,000 × *g*, washed with 50 mM EPPS (pH 8.3) and stored at -80°C until purification. 10 grams of cell paste were resuspended in 50 mM Na_2_HPO_4_/NaH_2_PO_4_ (pH 7.4), 300 mM NaCl (Resuspension Buffer) and lysed by microfluidizer (4 cycles at 30 kpsi). Lysed cells were centrifuged at 18,000 RPM and the supernatant fraction containing the soluble protein(s) of interest (POI) were taken for purification. The His-tagged proteins were purified using HisPur Ni-NTA resin (Thermo Fisher Scientific), at RT. Protein binding to the column was carried out in Resuspension Buffer containing 20 mM imidazole. The bound protein(s) was washed with increasing concentrations of imidazole (20 to 50 mM) in Resuspension Buffer and eluted in 10 column volumes of this buffer containing a gradient of 300-600 mM imidazole. Fractions containing the POI, as inferred by analysis with 12% SDS-PAGE stained with Coomassie Blue, were pooled and dialyzed against 50 mM EPPS (pH 8.0), 300 mM NaCl. Tagged proteins were processed with TEV protease to remove the respective N or C-terminal tags according to the methodology described in [42]. The cleaved protein mixture was subjected to a second round of purification by chromatography on Ni-NTA resin following the same gradient for elution. Flow through fractions containing tag less APRT1 or APRT2, as inferred by 12% SDS-PAGE stained with Coomassie Blue, were pooled and dialyzed against 50 mM HEPES (pH 7.4) 300mM NaCl, 50 mM L-Arginine, and 50 mM L-Glutamate. Homogeneous APRT1 was stored in the presence of 10% glycerol (v/v), 2.5mM TCEP at -80°C. Homogeneous APRT2 was stored in the presence of 10% glycerol (v/v) at -80°C.

### Discontinuous APRT activity assay (HPLC)

All chemicals were purchased from Sigma Aldrich, unless stated otherwise. PRPP concentrations were corrected for its purity as indicated by the supplier (75% purity). 9-deazaadenine was purchased from AA blocks (AA0037QI). Unless otherwise indicated, all kinetic assays were performed at RT.

For the forward (biosynthetic) assay of APRT, reaction mixtures contained 50 mM EPPS, 12 mM MgCl_2_, at pH 8.3 (Assay Buffer), with 1 mM PRPP, 150 μM adenine, 0.03 U inorganic pyrophosphatase (EC 3.6.1.1, Thermo Fisher Scientific) and APRT1 (0.1-25 μM) or APRT2 (0.2-25 μM). Reactions were incubated for 30 min and overnight, and were stopped by filtration to remove enzymes, using an Amicon Ultra (0.5 mL) filter with a 10,000 kDa cutoff, followed by centrifugation at 14,000 RPM for 25 min. Reverse reaction mixtures were treated similarly, and reactions mixtures containing 150 μM AMP, 1 mM inorganic pyrophosphate, and 4 μM *E. coli* K12 adenine deaminase (EC 3.5.4.2) [43], and APRT1 (0.1-25 μM) or APRT2 (0.2-25 μM). Individual chromatography standards contained 150 μM of adenine, AMP, or hypoxanthine in Assay Buffer. Negative controls were set in the same conditions as reaction mixtures, in the absence of APRT1 or APRT2. The nucleobase/nucleotide content of samples was analyzed by HPLC (high-performance liquid chromatography) using an Ultimate 3000 (Thermo Fisher Scientific) system equipped with a diode-array detector (DAD). Samples were analyzed on a Synergi Fusion-RP (4 μM, 4.6 × 150 mm) column. 15 μL of samples were injected onto the column and eluted with a gradient from 3 to 40% Solvent B over 10 minutes at flow rate of 1 mL min^-1^. Solvent A: 20mM ammonium acetate pH 4.5, Solvent B: 100% acetonitrile. Nucleobases and nucleotides were monitored at 260 nm.

### Continuous coupled enzyme assay for APRT activity

Steady-state kinetic assays for the forward reaction were measured using a coupled enzyme system composed of myokinase (MK), pyruvate kinase (PK), and lactate dehydrogenase (LDH) to catalyze the conversion of the AMP product to, respectively, ADP, pyruvate, lactate and NAD^+^; formation of the latter was monitored spectrophotometrically at 340nm (Scheme 1). The assays were carried out following the method described in [44] with some modifications. All assays were carried out in 100 mM HEPES buffer (pH 7.8) in the presence of 25 mM MgCl_2_, 1 mM ATP, 0.5 mM PEP, 300 μM NADH, 36 U mL^-1^ MK, 50 U mL^-1^ PK, and 36 U mL^-1^ LDH in a total volume of 250 μL. The conversion of NADH into NAD^+^ was monitored at 340 nm (ε = 6220 M^-1^ cm^-1^).

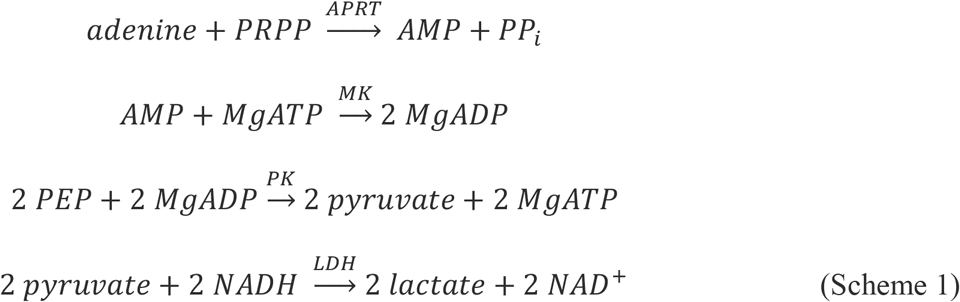

Apparent kinetic parameters for APRT1 and APRT2 (app *K*_*M*_, app *k*_*cat*_, and app *k*_*cat*_*/K*_*M*_) were determined for adenine from initial velocity measurements in duplicate or triplicate with at least seven varied substrate concentrations, and at apparent saturating concentrations of PRPP (1 mM). Reaction mixtures of 250 μL contained adenine 2.5-60 μM in the presence of apparent saturating concentrations of PRPP (1 mM). Kinetic parameters were calculated by fitting of initial rate data to Eq (1) using GraphPad Prism 9.2 software [45]. Initial velocity data (*K*_*M*_, *k*_*cat*_, and *k*_*cat*_*/K*_*M*_) for recombinant APRT1 (200 nM) were determined as plots of changing-fixed concentration of adenine (1 μM -20 μM) *vs*. variable PRPP (10 – 250 μM), or as changing-fixed concentrations of PRPP (25 μM -150 μM) *vs*. variable adenine (1.5 – 50 μM), and data were fitted to Eq (2).

#### Initial velocity data using mixtures of APRT1 and APRT2

Purified APRT1 (2.5 μM) and APRT2 (2.5 μM) were incubated in a 1:1 equimolar ratio on ice for 4 hours. Kinetic assays were performed as described above for the continuous APRT activity assay. 200 nM of APRT (APRT1 or the APRT1:APRT2 mixture) was added to the assay mixture to start the reaction. Assays were performed in triplicate.

### Inhibition Studies

To determine if the inhibition pattern of 9-deazaadenine against adenine and PRPP was competitive, uncompetitive, or noncompetitive, initial velocity data were collected in the presence of five apparent non-saturating concentrations of 9-deazaadenine. An inhibition pattern of 9-deazaadenine *vs*. adenine was ascertained by initial velocity data plotted as double-reciprocal plots of changing-fixed concentration of 9-deazaadenine (0 μM -300 μM) *vs*. variable adenine (2.5 – 150 μM) at apparent saturating conditions of PRPP (1 mM). The 9-deazaadenine inhibition pattern against PRPP was determined as double reciprocal plots of initial velocity data in the presence of changing-fixed concentrations of 9-deazaadenine (0 μM -600 μM) *vs*. variable PRPP (12.5 – 1000 μM) at apparent saturating concentration of adenine (100 μM). Inhibition data were fitted to Eq (3)-(5). AMP formation was measured as previously described with the continuous coupled-enzyme APRT activity assay.

### Crosslinking assays

Homogeneous APRT1, APRT2-Ntag, and APRT2-Ctag were treated with dimethyl suberimidate (DMS) according to the method of Davies and Stark [46], in order to assess the extent to which homodimers or other multimeric forms existed in solution, as exemplified by protein cross-linking. Briefly, reaction mixtures containing 0.2 M triethanolamine at pH 8.5, and purified APRT (17.8, 35, 69, or 140 μg mL^-1^) in the presence or absence of 8.7 mM DMS were incubated at RT for 5-8 hours, then denatured with SDS-PAGE sample buffer. The samples were subjected to analysis by 12% SDS-PAGE stained with Coomassie Blue to ascertain the respective levels of monomeric and multimeric forms of the enzymes that were present.

### Evaluation of other nucleobase substrates of APRT2

APRT2 enzyme activity was tested spectrophotometrically using alternative N-containing nucleophiles as possible substrates, including orotate, uracil, hypoxanthine, guanine, and xanthine. All assays were conducted in assay buffer containing 1 mM PRPP, APRT2 (25 μM to 50 μM), and 150 μM of base, in a final volume of 250 μL. Possible activities were assayed as follows: orotate phosphoribosyltransferase (OPRT – EC 2.4.2.10) activity was measured as described in [47], where formation of OMP was monitored at 293 nm. Hypoxanthine-guanine-xanthine phosphoribosyltransferase (HGXPRT – EC 2.4.2.8) activity was measured as described in [48, 49]. Formation of IMP, GMP, and XMP were monitored at, respectively, 245 nm, 257.5 nm, and 255 nm. Uracil phosphoribosyltransferase (UPRT – EC 2.4.2.9) activity was measured as described in [50], in which monitoring of UMP formation was conducted at 280 nm. The ability of APRT2 to catalyze the synthesis of PRPP from ATP and ribose-5-phosphate (PRPP synthetase activity -EC 2.7.6.1) was evaluated using the described coupled-enzyme assay in which the product AMP is measured by coupling the PRPP synthetase reaction 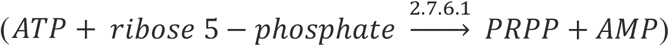 to MK, PK, and LDH (Scheme 1) [51, 52] in reaction mixtures containing 100 mM HEPES (pH 7.8), 25 mM MgCl_2_, 2 mM ATP, 0.5 mM PEP, 300 μM NADH, 150 μM ribose-5-phosphate, 36 U mL^-1^ MK, 50 U mL^-1^ PK, and 36 U mL^-1^ LDH in a total volume of 250 μL. The ability of APRT2 to catalyze the reactions of adenosine phosphorylase (EC 2.4.2.1), and 5’-nucleotidase (EC 3.1.3.5) activities were also analyzed using the same methodology described for the discontinuous APRT activity assay (HPLC) above, with the following modifications. Samples (15 μL) were injected onto the column and eluted with a gradient from 5 -40% Solvent B over 15.1 minutes at a flow rate of 0.425 mL min^-1^. Solvent A: 20 mM ammonium acetate pH 4.5, Solvent B: 100% acetonitrile. Nucleobases and nucleosides were monitored at 260 nm. Assays to detect adenosine phosphorylase (EC 2.4.2.1) activity were carried out in assay buffer and contained 1 mM PRPP, 150 μM adenosine, 0.03 U inorganic pyrophosphatase (EC 3.6.1.1, Thermo Fisher Scientific) and APRT2 (25 or 50 μM). Assays to detect 5’-nucleotidase (EC 3.1.3.5) activity were carried out in assay buffer and contained 150 μM AMP, 4 μM *E. coli* K12 adenine deaminase (EC 3.5.4.2) [43], and APRT2 (25 or 50 μM). Assays were incubated for 30 min and overnight. Reactions were stopped by filtration to remove enzymes, using an Amicon Ultra (0.5 mL) filter with a 10,000 kDa cutoff, followed by centrifugation at 14,000 RPM for 25 min.

### Cyanogen bromide cleavage of proteins

Cyanogen bromide cleavage reactions were performed as described in [53]. 10 μL of 5 M HCl were used to acidify 250 μg/mL of APRT2-Ntag, APRT2-Ctag, or APRT Ec (*E. coli* expressed APRT2 stored in 5 mM DTT) in a total volume of 95 μL (Milli-Q water). 5 M Cyanogen bromide was added in 200 molar excess (200:1, CNBr/Met residue). APRT2 contains 5 Met residues, approximately 2.1% Met residue content. Samples were vortexed, covered with aluminum foil, and incubated for 24 hours at RT. The reaction was stopped by adding 5 volumes of Milli-Q water. Samples were dried using a vacufuge (60°C, 2 hours), resuspended in 50 μL water, and analyzed by 12% SDS-PAGE stained with Coomassie Blue.

### Native Mass Spectrometry (MS)

APRT2-Ntag and APRT2-Ctag were buffer exchanged into a volatile buffer (200 mM ammonium acetate, pH 7.4) using a Micro Bio-Spin 6 device (Bio-Rad). Samples were loaded into pulled glass capillaries prepared in house, as described in [54], and introduced into an Orbitrap mass spectrometer (Exactive Plus with extended mass range (EMR), Thermo Scientific) by means of electro-spray ionization. The instrumental parameters were tuned to preserve the native-like structure of the proteins as described in [54].

### Cloning, expression, and purification of wild-type and mutants of APRT2

APRT2 wild-type (WT), and mutants M73Q, and M128Q were codon optimized for expression in *E. coli* and cloned into pET-28a(+) vector, using *Nde*I and *Hind*III restriction sites (Genscript, USA). Optimal expression of all three proteins was obtained in BL21(DE3) cells, using Terrific-Broth media.

The plasmids pET-28a(+)::*aprt2*, pET-28a(+)::*aprt2M73Q* and pET-28a(+)::*aprt2M128Q* were individually transformed into BL21(DE3) cells by heat-shock, and plates containing kanamycin 50 μg mL^-1^ were incubated overnight at 37°C. A single colony of each strain was used to inoculate 50 mL cultures of Terrific-Broth media in the presence of kanamycin 50 μg mL^-1^. Starting cultures were incubated overnight at 37°C in an orbital shaker at 180 RPM. Starting culture (13mL) was used to inoculate 500 mL of Terrific-Broth media in the presence of kanamycin 50 μg mL^-1^. Cultures were grown at 37°C and 180 RPM until the O.D._600_ reached 0.4-0.6. Protein expression was induced by the addition of IPTG to a final concentration of 1mM. Cells were further grown under these conditions for 24 hours. Cells were harvested by centrifugation at 8,000 × *g*, washed with 50 mM Tris HCl, 300 mM NaCl (pH 7.4) and stored at -20°C until purification. 5 grams of cell paste were resuspended in Resuspension Buffer and disrupted by sonication (15 cycles of 10 seconds at 60% amplitude) in the presence of 0.2 mg mL^-1^ lysozyme. After centrifugation at 48,000 × *g*, the N-terminal His-tagged protein was purified in a single step protocol using HisPur Ni-NTA resin (Thermo Fisher Scientific), equilibrated at RT. Protein binding was carried in Resuspension Buffer supplemented with 20 mM imidazole. The bound protein was washed with increasing concentrations of imidazole (20 to 50 mM) in Resuspension Buffer and eluted in 10 column volumes of Resuspension Buffer containing 300-600 mM imidazole. Fractions containing apparently homogeneous APRT2 WT, APRT2 M73Q, and APRT2 M128Q, as inferred by SDS-PAGE (12% polyacrylamide; stained with Coomassie Blue), were pooled and dialyzed against 50mM EPPS, 300mM NaCl, (pH 8.0). 5 mM DTT was added to APRT2 WT to ensure Met residues potentially oxidized during cell lysis were fully reduced. Homogeneous recombinant proteins were stored in presence of 10% (v/v) glycerol at -80°C.

APRT2 mutant M156Q was cloned, expressed, and purified as described above, with the following modifications: APRT2 M156Q best expression was observed in C43(DE3) cells, using Terrific-Broth media. After IPTG induction, cells were grown for 24 hours at 18°C. Homogeneous APRT2 M156Q, as inferred by 12% SDS-PAGE stained with Coomassie Blue, was pooled, dialyzed, and stored under the same conditions described above.

### Data Analysis

Data analysis was performed using GraphPad Prism 9.2 software. Initial velocity data obtained at variable concentrations of a single varied substrate *A* at fixed concentrations of another substrate were fitted to Eq (1), in which *v* is the initial velocity, E_t_ is the enzyme concentration, *k*_*cat*_ is the turnover number, *A* is the concentration of the variable substrate, and *K*_*a*_ is the apparent Michaelis constant.

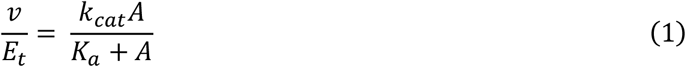

Initial velocity data obtained at variable concentrations of a single substrate *A* at changing-concentrations of a second substrate *B* were fitted to Eq (2), in which *K*_*ia*_ and *K*_*a*_ are the respective dissociation and Michaelis constants of *A*, and *K*_*b*_ is the Michaelis constant of substrate *B*.

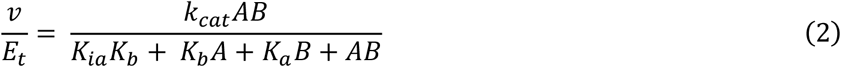

For inhibition studies in which double-reciprocal plots of inhibitor (*I*) *vs*. variable substrate (*A*) conformed to apparent competitive, noncompetitive, or uncompetitive inhibition, respectively, initial velocity data (*v*) were fitted to Eq (3)-(5), for which E_t_ is the concentration of enzyme, *k*_*cat*_ and *K*_*a*_ are, respectively, the turnover number and the Michaelis constant for substrate A, and *K*_*is*_ and *K*_*ii*_ are respectively, the apparent slope and intercept inhibition constants.

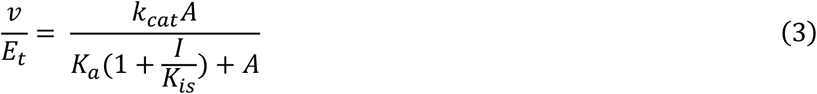

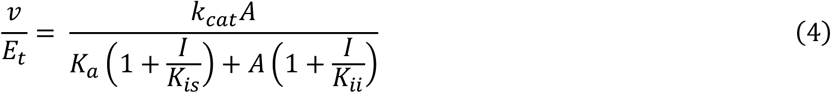

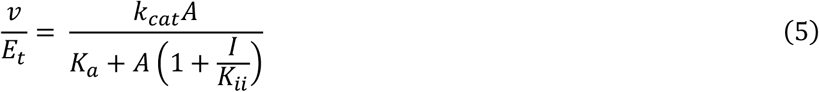

## Results and Discussion

*T. brucei brucei* genome data includes two putative *aprt* genes (APRT1 – Tb927.7.1780, and APRT2 – Tb927.7.1790) located in tandem on chromosome 7. To ascertain the catalytic activity of the protein products of these two genes, both were cloned, expressed, and purified individually. Upon evaluation of *in vitro* activity, APRT1 was capable of catalyzing the forward and reverse reactions of APRT (Fig 1, S4 Fig, and S5 Fig), while APRT2 catalysis of the forward APRT reaction is negligible (Table 1). From the steady-state kinetic analysis of the forward reactions catalyzed by APRT1 and APRT2 at apparent saturating fixed concentration of PRPP, we determined apparent *k*_*cat*_ and *K*_*M*_ values for adenine (Table 1). APRT2 displays a significantly lower *k*_*cat*_*/K*_*M*_ value (> 40,000-fold decrease) when compared to APRT1 (Table 1). Although the reverse activity catalyzed by APRT1 was detected, the extent of the conversion of AMP into adenine is minor when compared to APRT1 forward activity (2.5% conversion in an assay containing 25μM APRT1 at RT, 30-minute incubation S4 Fig). This observation supports the assumption that the primary cellular role of APRT1 is scavenging 6-amino purine bases from hosts and their incorporation into nucleotides and nucleic acids via the purine salvage pathway. A greater preference for the forward reaction can be expected for obligatory purine auxotrophs.

**Table 1.**
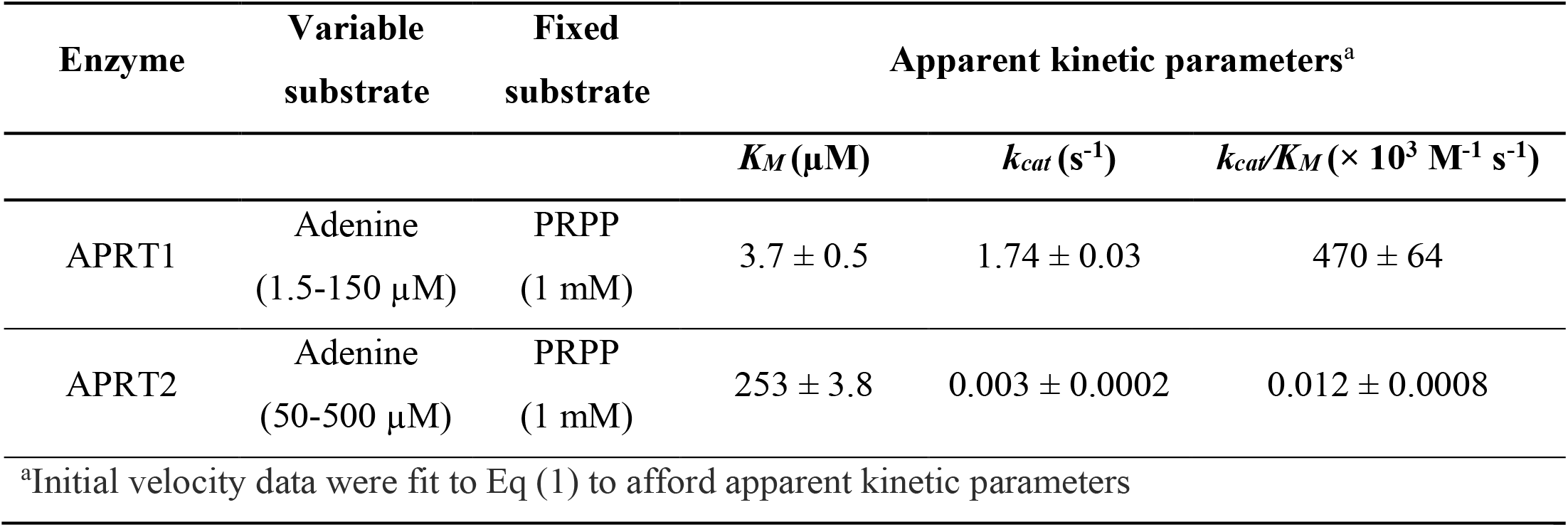
Apparent kinetic parameters of APRT1 and APRT2.

Notably, catalysis of the APRT biosynthetic reaction by APRT2 is negligible when compared to APRT1. Endogenous adenine is present in human plasma at a concentration of approximately 9 nM [55], a value almost 30,000 lower than the APRT2 determined apparent *K*_*M*_ for adenine (Table 1), implying the APRT activity detected is not biologically relevant under normal metabolic conditions. Adenine concentration in human serum is tightly regulated, as higher adenine concentrations are related to numerous medical conditions [56-58]. Moreover, high concentrations of adenine are toxic to bloodstream stage and procyclic stage *T. brucei* (IC_50_ ∼ 300 μM) [35], corroborating our findings that APRT2 is not a physiologically relevant APRT in *T. brucei*. Furthermore, the nearly 100-fold increase in apparent *K*_*M*_ of adenine for APRT2, when compared to APRT1, suggests adenine is not the natural substrate for APRT2 (Table 1).

APRT1 was further characterized in terms of its kinetic mechanism. The initial velocity data of PRPP *vs*. adenine when plotted in double-reciprocal form (S5 Fig, S6 Fig) conformed to an intersecting pattern, which indicated that a sequential bi-substrate mechanism is operative for APRT1. Fitting of the data to Eq (2) afforded the kinetic parameters shown in Table 2. In kind, double-reciprocal plots for the initial velocity data of adenine *vs*. PRPP also conformed to an intersecting pattern, providing nearly identical kinetic parameters for both PRPP and adenine, when compared to the PRPP *vs*. adenine initial velocity data (S5 Fig). *Leishmania donovani*, also a unicellular parasitic protozoa purine auxotroph with complex life cycle encompassing insect and human hosts, has a single cytosolic APRT (*Ld*APRT). When compared to *Ld*APRT, APRT1 presents a similar *K*_*M*_ for adenine but displays a decreased catalytic efficiency (*k*_*cat*_*/K*_*M*_) for both substrates, with a nearly 30-fold decrease for adenine, and a 10-fold decrease for PRPP (Table 2). Similarly, the protozoan parasite *Giardia lamblia* also contains a single APRT (*Gl*APRT), and when compared to APRT1 presents similar *k*_*cat*_*/K*_*M*_ values for both PRPP and adenine.

**Table 2.** Initial velocity data for the biosynthetic reaction of APRT1 and other characterized parasitic protozoan APRTs.

| Enzyme | Variable<br>Substrate | Kinetic parameters<sup>a</sup> |  |  |  |  |  |
| --- | --- | --- | --- | --- | --- | --- | --- |
| | | $K_{PRPP}$<br>($\mu\text{M}$) | $K_{i,PRPP}$<br>($\mu\text{M}$) | $k_{cat}/K_{PRPP}$<br>($\times 10^3 \text{ M}^{-1}\text{s}^{-1}$) | $K_{Ade}$<br>($\mu\text{M}$) | $k_{cat}/K_{Ade}$<br>($\times 10^3 \text{ M}^{-1}\text{s}^{-1}$) | $k_{cat}$<br>($\text{s}^{-1}$) |
| *TbbAPRT1* | Adenine<br>(1.5-50) | $77 \pm 12$ | $61 \pm 13$ | $8.1 \pm 1.3$ | $2.2 \pm 0.7$ | $282 \pm 90$ | $0.62 \pm 0.02$ |
| | PRPP<br>(10-250) | $68 \pm 11$ | $22 \pm 13$ | $9.0 \pm 1.6$ | $2.3 \pm 0.4$ | $265 \pm 49$ | $0.61 \pm 0.04$ |
| *LdAPRT*<sup>b</sup> | Adenine<br>(2.5 –50) | $25 \pm 6$ | N.D. | $708 \pm 282$ | $2.3 \pm 1.3$ | $7782 \pm 5027$ | $17.9 \pm 5.6$ |
| *GlAPRT*<sup>c</sup> | Adenine<br>(3–10) | $143 \pm 47$ | 376 | $19.6 \pm 6.5$ | $4.2 \pm 0.6$ | $595 \pm 91$ | $2.5 \pm 0.14$ |

<sup>a</sup>Initial velocity data were fit to Eq (2) to afford kinetic parameters. N.D. = not determined
<sup>b</sup>Data for *Leishmania donovani* APRT (*LdAPRT*) obtained by Bashor *et. al.*, 2002
<sup>c</sup>Data for *Giardia lamblia* APRT (*GlAPRT*) obtained by Sarver *et. al.*, 2002

The order of substrate binding and product release for APRT1 was determined by analysis of the dead-end inhibitor 9-deazaadenine (Table 3, Fig 2, S7 Fig), where 9-deazaadenine behaves as a competitive inhibitor to adenine as indicated by the double-reciprocal plots of initial velocity data at apparent saturating concentrations of PRPP, fixed-changed concentrations of 9-deazaadenine, and variable concentrations of adenine (Fig 2). At increasing fixed concentrations of 9-deazaadenine the intercept is not changed (1/*V*_*max*_) but the slope (*K*_*M*_*/V*_*max*_) is changed, indicative of competitive inhibition and suggesting 9-deazaadenine and adenine bind to the same enzyme forms.

**Table 3.**
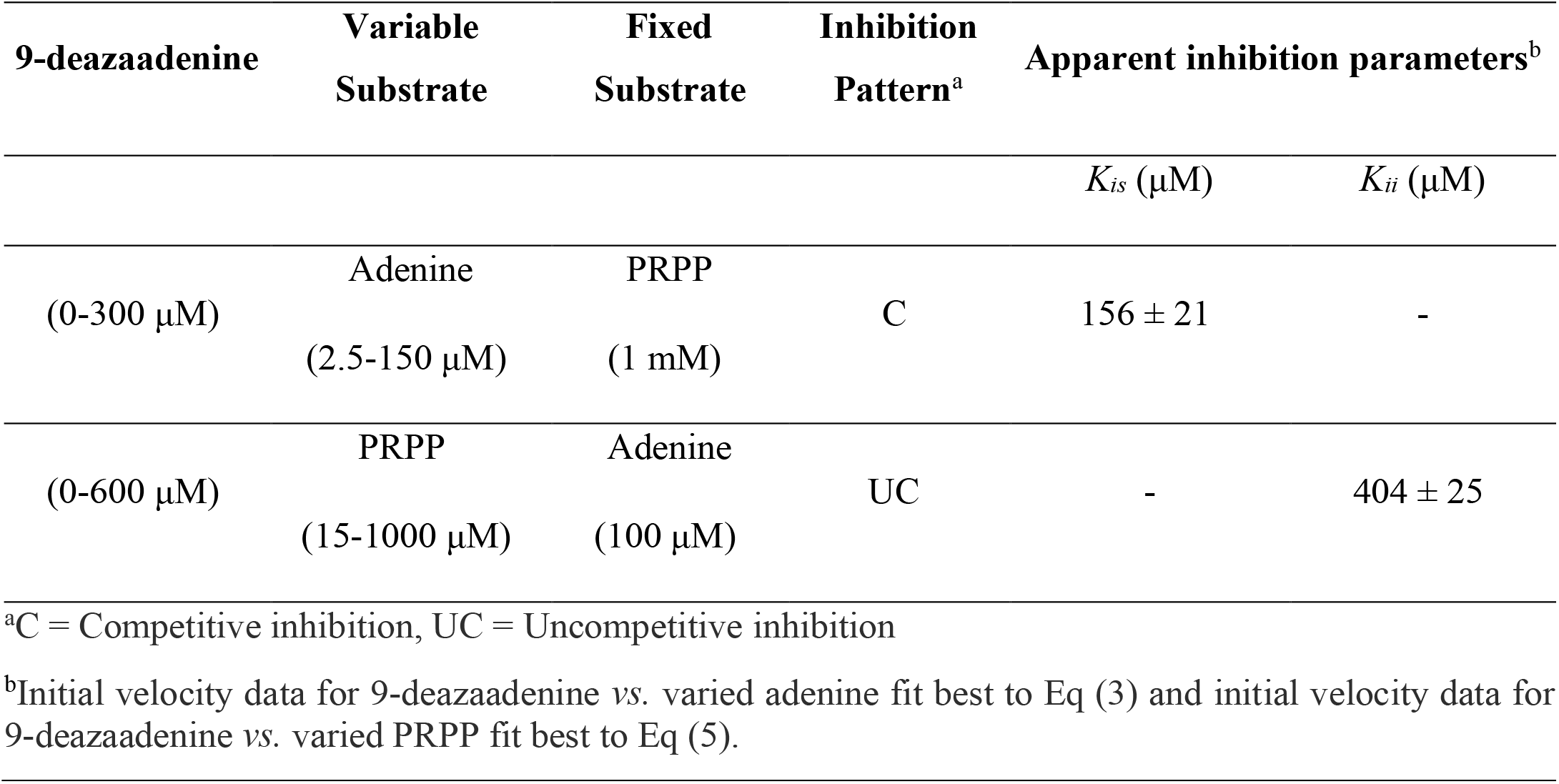
Dead-End inhibition data of 9-deazaadenine against varied adenine and PRPP.

**Fig 2.**
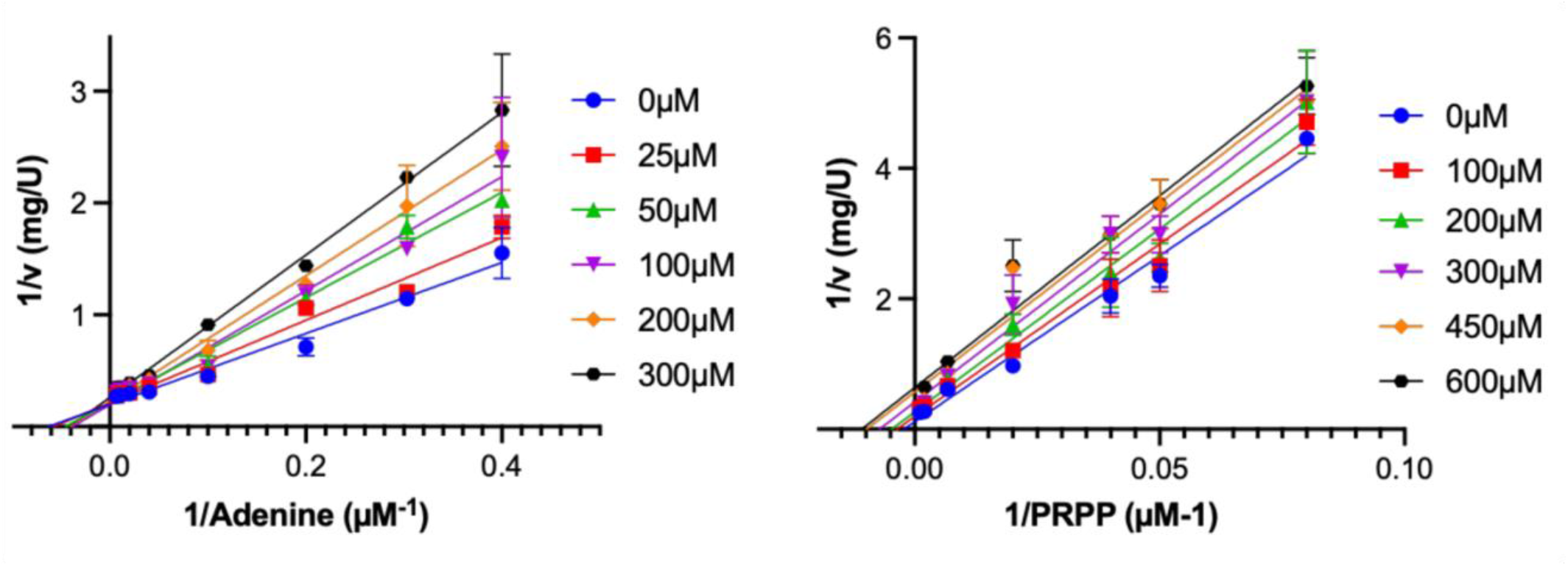
Double reciprocal plots of initial velocity data in the presence of the dead-end inhibitor 9-deazaadenine. **(A)** 9-deazaadenine is a competitive inhibitor *vs*. varied adenine (2.5-150 μM) at fixed apparent saturating concentrations of PRPP (1 mM) **(B)** 9-deazaadenine is an uncompetitive inhibitor *vs*. varied PRPP (12.5-1000 μM) at fixed apparent saturating concentrations of adenine (100 μM).

Double reciprocal plots of the initial velocity data at apparent saturating concentrations of adenine (100 μM) at fixed-changed concentrations of 9-deazaadenine *vs*. varying PRPP provided an apparent uncompetitive pattern. At increasing concentrations of 9-deazaadenine the intercept is changed (1/*V*_*max*_) but the slope (*K*_*M*_*/V*_*max*_) is unaffected, suggesting that PRPP binds to the enzyme before the addition of 9-deazaadenine. These results agree with the initial velocity data for APRT1, (Table 1 and Table 2) and reveal a Bi-Bi sequential ordered kinetic mechanism, where the phosphoribosyl donor, PRPP, binds to free enzyme, followed by the second substrate, adenine. After catalysis, PP_i_ is the first product to dissociate from the enzyme-products complex, followed by the nucleotide monophosphate, AMP (S8 Fig).

An ordered sequential mechanism has been previously described for *Ld*APRT [59]. The cytosolic *Ld*APRT [56-58] has higher sequence conservation to the cytosolic *T. brucei brucei* [35, 41] APRT1 (52%) than to the glycosomal APRT2 (27%) [35, 41]. Interestingly, *Gl*APRT was described to follow a unique reaction mechanism among phosphoribosyltransferases, in which there is a random addition of substrates (PRPP and adenine) followed by catalysis and an ordered release of the products, where PPi is the first product to leave the enzyme-products complex followed by AMP. *Gl*APRT primary sequence has 35% identity to APRT1 and 29% identity to APRT2.

APRT1 has been reported to adopt a dimeric quaternary structure (PDB code 5VN4), so we investigated whether a heterodimer of APRT1:APRT2 could form *in vitro* and provide a hybrid enzyme capable of catalysis. AMP formation was evaluated in reaction mixtures containing equimolar amounts of the two APRTs, which could give rise to three sets of dimers: APRT1 homodimers, APRT2 homodimers, and APRT1:APRT2 heterodimers in various ratios. AMP formation was reduced 2-fold from a control reaction mixture containing only APRT1 (Fig 3). The reduction in APRT activity is explained by the 1:1 dilution of APRT1, thereby indicating a catalytically active APRT1:APRT2 heterodimer is not formed.

**Fig 3.**
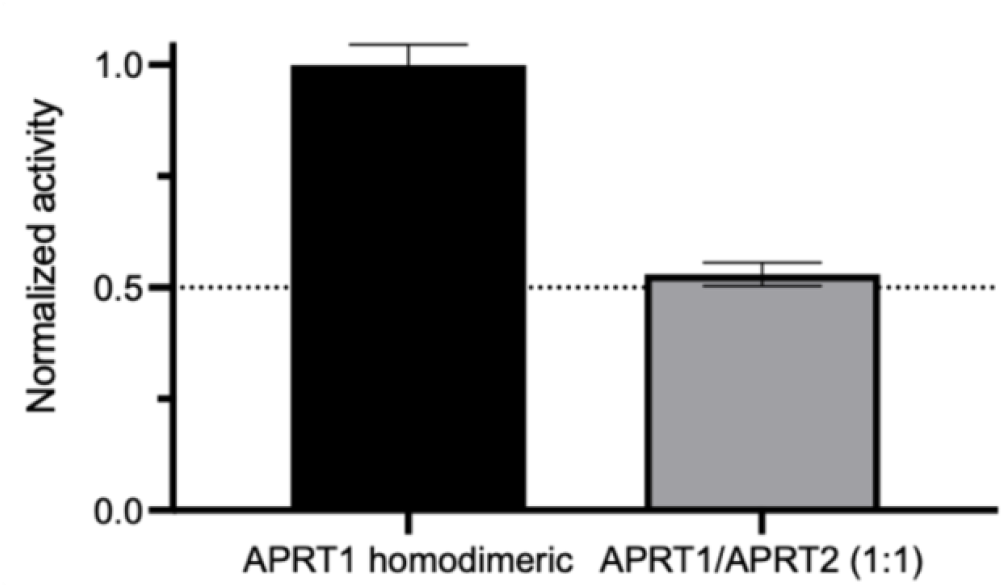
APRT activity as measured by AMP formation of reactions mixtures composed of 150 μM adenine, 1 mM PRPP and 200 nM APRT1 (left) or 200 nM APRT mixture (100 nM APRT1: 100 nM APRT2, right).

A recent study revealed APRT1 and APRT2 do not interact *in vivo*, with APRT1 displaying homodimeric properties both *in vivo* and *in vitro* [41], in agreement to results showed in Fig 3. The quaternary structure of APRT1 was evaluated *in vitro* by treating APRT1 and APRT2 with DMS, an agent that chemically crosslinks two lysine residues within a monomer and also between lysine residues found in two or more associated monomers (Fig 4) [46]. Analysis of crosslinked proteins by denaturing polyacrylamide electrophoresis will show homodimers and other multimers, indicating quaternary structure, which will still be observed after significant protein dilution. Purified APRT1 (17.8 – 140 μg/mL) was treated with 8.7 mM DMS at pH 8.5. Our results corroborated the finding that APRT1 forms homodimers *in vitro*, as indicated by the observed dimeric structures, which is maintained after 4-fold dilution. To ensure the C-terminally localized glycosomal signal sequence of APRT2 did not obstruct dimerization events, APRT2 crosslinking assays were performed using recombinant enzyme expressed using pPICZ-Ntag (APRT2-Ntag) and pPICZ-Ctag (APRT2-Ctag) vectors (S2 Fig). There was no significant difference observed between APRT2-Ntag and APRT2-Ctag, and both show less frequent formation of a dimeric quaternary structure, which is not maintained after dilution as compared to APRT1 under identical concentrations and conditions (Fig 4).

**Fig 4.**
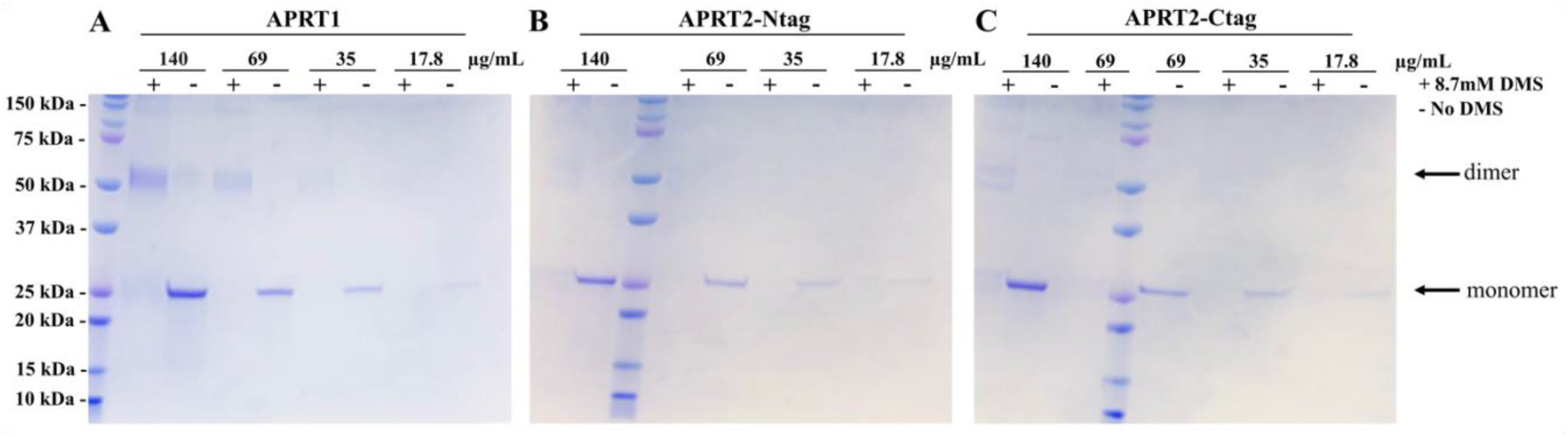
SDS-PAGE analysis of chemically crosslinked recombinant APRT1 and APRT2. APRT1 (**A**) homodimeric quaternary structure can be visualized under crosslinking assay conditions in concentrations ≥ 35 μg mL^-1^, in presence of crosslink reagent DMS (8.7 mM), indicated by (+). Negative control lanes (-) show APRT1 monomers. APRT2 expressed with N-terminal (**B**) or C-terminal (**C**) tags.

APRTs belong to the type 1 phosphoribosyltransferases (Type 1 PRTases) family of homologous enzymes and regulatory proteins of the nucleotide synthesis and salvage pathways. The alignment of APRT1 and APRT2 primary sequences using ClustalW [60] showed that only 22% of their amino acids are conserved (Fig 5). Type 1 PRTases share highly conserved structures, but low levels of sequence identity except for the PRPP binding motif [61]. This motif is conserved in both APRT1 (VVLIDDVIATGGT), and APRT2 (VLIVDDFIGTGST) (Fig 5, red underline). A BLASTp search (https://blast.ncbi.nlm.nih.gov/Blast.cgi) search using APRT2 as the query sequence classifies it as a Type 1 PRTase, with no other homology matches. The lack of relevant APRT activity suggests that APRT2 may play an alternative metabolic role. Low sequence identity does not preclude conservation of function for this family of proteins and since APRT2 presents the conserved PRPP binding motif, we tested the putative APRT2 activity as a Type 1 PRTase using other potential substrates. No phosphoribosyl transfer could be detected in the presence of hypoxanthine, guanine, xanthine, orotate, or uracil, ruling out possible HGXPRT (EC 2.4.2.8), OPRT (EC 2.4.2.10), or UPRT (EC 2.4.2.9) activities respectively (S9 Fig). Neither adenosine phosphorylase (EC 2.4.2.1), 5’-nucleotidase (EC 3.1.3.5), nor PRPP synthetase (EC 2.7.6.1) activities could be detected under our assay conditions (S9 Fig and S10 Fig).

**Fig 5.**
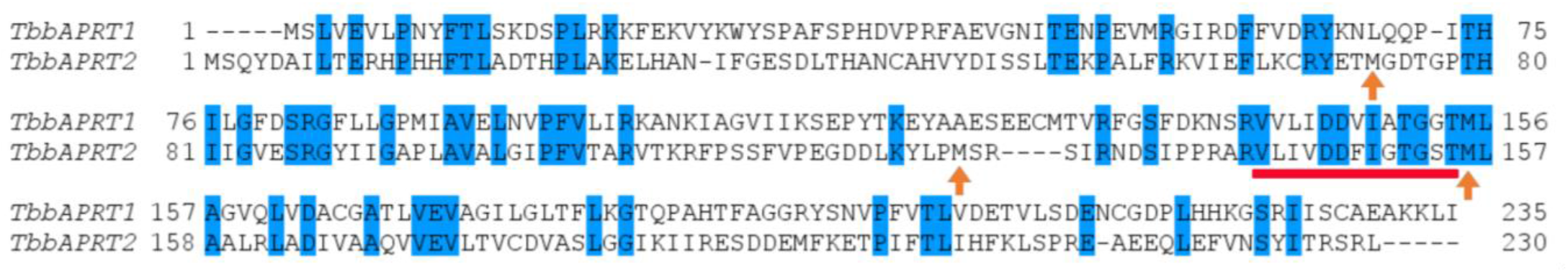
Alignment of *T. b. brucei* APRT1 and APRT2. Conserved residues are shaded in blue. The conserved PRPP binding domain - a 13 residues-long active site loop - is underlined in red. APRT2 mutated methionine residues (M to Q) are indicated by orange arrows.

A recent publication showed that RNAi silencing of both APRT1 and APRT2 activity in *T. brucei* bloodstream stage cells did not produce a major growth phenotype, indicating the HGXPRT activity is adequate to maintain cell growth under normal growth conditions [41]. When the same RNAi cell line was tested as bloodstream stage cells grown in medium containing adenine as the sole available purine, a significant growth phenotype was observed suggesting the APRTs are the only enzymes that can incorporate adenine into the purine salvage pathway. The same publication reported another RNAi cell line, silencing APRT1 activity only, which displayed a growth phenotype similar to the bloodstream stage cells grown in medium containing only adenine, suggesting APRT2 is not able to compensate for the loss of APRT1 even when media contained up to 50 μM adenine [41].

What thus, can be the role or function of APRT2? The existing literature on APRT2 provides no data on its catalysis or metabolic role. Along with its localization to the glycosome and expression in the procyclic life stage [35, 62], a co-immunoprecipitation study identified APRT2 as a protein associated with TbTim17, the major component of the mitochondrial inner membrane translocase complex in trypanosomes, with detection of a post-translational modification (PTM), in which a methionine residue of APRT2 was fully oxidized to methionine sulfoxide (MetO) [40]. PTMs have been described in trypanosomes [63] and are a freely reversible mechanism of control at the protein level that would enable the parasites to react quickly to environmental changes, including the presumably abrupt changes associated with vector (tsetse fly) to host transmission [64]. The metabolic change in procyclic trypanosomes from glycolysis to oxidative phosphorylation due to activation of the mitochondrion [65, 66] could enable reactive oxygen species (ROS) to oxidize a Met residue to MetO [67]. The reversible formation of MetO by ROS as a mechanism of enzymatic activation has been described [68, 69]; and the enzyme that mediates the reversal of this oxidation, methionine sulfoxide reductase, is putatively localized in the *T. brucei* genome (Tb927.8.550).

Utilizing a eukaryotic expression, such as *P. pastoris*, provided many advantages including occurrence of PTMs [70]. PTMs such as the oxidation of methionine residues have been previously reported in other heterologous proteins expressed in *P. pastoris* [71, 72]. We employed the methionine-specific agent, cyanogen bromide (CNBr), to determine if (any) methionine residues in the APRT2 expressed in *P. pastoris* were oxidized into MetO. We utilized APRT2 expressed in *E. coli* (APRT2 Ec) - an expression system incapable of performing PTMs - as a negative control. APRT2 Ec was stored in the presence of the reducing agent DTT (5 mM) following purification to ensure methionine residues potentially oxidized during cell lysis were fully reduced. Peptide cleavage at the C-terminus of methionine residues in proteins by treatment with CNBr is a selective method to identify the presence of oxidized methionine residues in proteins as these will be refractory to this chemistry [73]. APRT2 has five methionine residues in its primary sequence. Treatment of recombinant APRT2 with 0.5 M HCl (final concentration) was followed by the addition of 250 mM CNBr. Analysis of the resulting protein fragments was observed utilizing SDS-PAGE. Recombinant APRT2 expressed in *P. pastoris* (APRT2-Ntag and APRT2-Ctag) provided identical patterns to the control (APRT2 Ec). Thereby indicating that both APRT2-Ntag and APRT2-Ctag, did not present a MetO PTM on any of the five known methionine residues present on its primary sequence (Fig 6 A). Indeed, the resulting protein fragments provide bands with apparent molecular masses indicated on S11 Fig, where none of the 5 methionine residues are oxidized to MetO. These results were confirmed with Native-MS, where a change of 16 Da on the protein subunit mass – equivalent to a MetO PTM, could not be identified (Fig 6 B and C). The native MS of recombinant APRT2-Ntag and APRT2-Ctag provided the measured masses of 25,860 and 26,637 Da, respectively. The theoretical monoisotopic mass of APRT2-Ntag and APRT2-Ctag are 25, 853 and 26,647 Da, respectively. The absence of a 16 Da increase from the theoretical monoisotopic mass of APRT2 monomers confirmed the lack of MetO formation. These results agreed with our analysis of the CNBr treatment, indicating that none of the recombinant APRT2s contained a MetO modification. Interestingly, the APRT2-Ntag also displayed a dimer with a mass of 51,739 Da, providing evidence that the C-terminal glycosomal signaling peptide may be involved in dimer formation. However, crosslinking recombinant APRT2-Ntag with DMS did not provide evidence for a dimeric quaternary structure, as the dimeric multimer is not maintained after sufficient dilution under the tested conditions, indicating APRT2-Ntag dimers are likely unstable (Fig 4).

**Fig 6.**
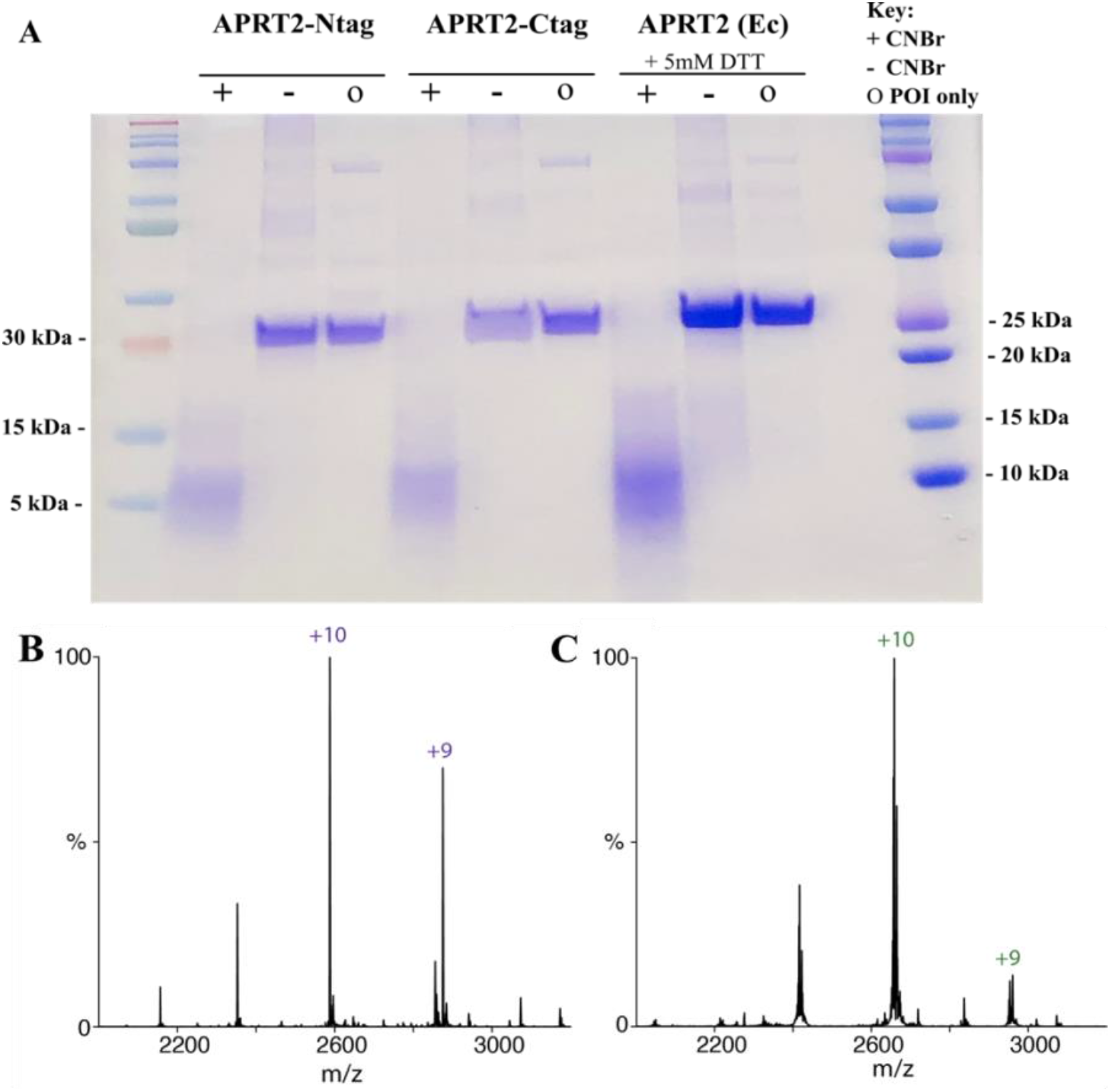
SDS-PAGE analysis of CNBr cleavage reaction (A) and Native-MS of recombinant APRT2s (B, C). **(A)** When APRT2 methionine residues are not oxidized to MetO, bands of molecular mass with the expected sizes indicated in S11 Fig would be observed. APRT2 expressed in *E. coli* and stored in the presence of 5 mM DTT (APRT2 Ec) was used as a control. The bands resulting from CNBr cleavage for the APRT2-Ntag, APRT2-Ctag, and APRT2 Ec are shown in the (+) lanes. Negative controls (-) and untreated native protein (o) show the band corresponding to the monomer mass. Native-MS of recombinant APRT2-Ntag **(B)** and APRT2-Ctag **(C)** provided the measured masses of 25,860 and 26,637 Da, respectively.

Due to the apparent absence of MetO formation in recombinant APRT2, we introduced methionine to glutamine (M to Q) mutations to mimic this PTM. The mutation of methionine residues to glutamine has previously been found to mimic the functional effects of a MetO, since both glutamine and MetO present an oxygen atom at the same position in their side chains and exhibit roughly the same hydrophobicity index value [67, 74]. The crystallographic structure of APRT1 shows two methionine residues located near the PRPP binding site and one methionine residue located near the subunit interface, respectively M129, M155, and M89, numbered according to APRT1 primary sequence (S12 Fig). The APRT2 methionine residues located in the corresponding positions to APRT1 (M73, M128, and M156, according to APRT2 the primary sequence – Fig 5, orange arrows) were mutated to glutamine and the recombinant mutated proteins were analyzed for APRT activity. The forward reaction was monitored in presence of inorganic pyrophosphatase (EC 3.6.1.1) to displace the reaction equilibrium towards AMP formation: 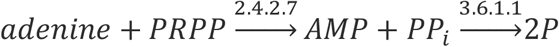. APRT1 was used as a reaction control, and the complete conversion of adenine into AMP was observed in both control assay conditions (30 minute and overnight incubation). The three APRT2 M to Q mutants (M73Q, M128Q, and M156Q) displayed negligible APRT activity even when the assay was performed overnight in the presence of high concentrations of enzyme (25 μM) (Fig 7, S13 Fig). These data suggested that the presence of a MetO does not increase nor elicit the catalytic APRT activity of APRT2, and the absence of APRT activity observed is indeed a reflection of a misannotation of Tb927.7.1790 as a putative APRT enzyme.

**Fig 7.**
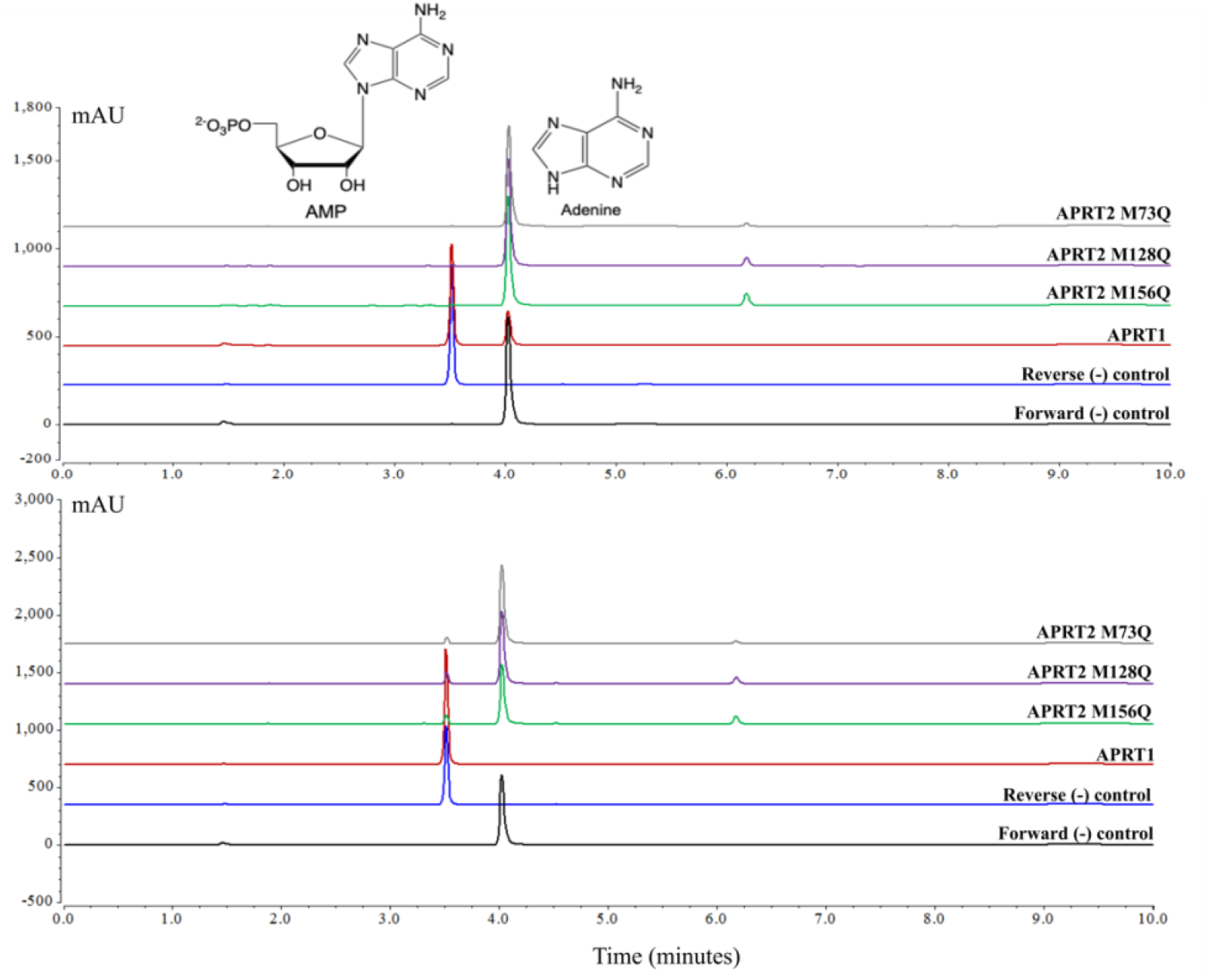
HPLC chromatograms of APRT2 M to Q mutants’ activity. Activity assays of the APRT2 M to Q mutants’ APRT activity assays with a 30 min (top graph) or overnight (bottom graph) incubation. Controls for the forward reaction showed the elution peak corresponding to adenine (Retention time (R_t_) = 4.2 minutes, black lines), and reverse reaction showed the elution peak corresponding to AMP (R_t_ = 3.6 minutes, blue lines). The eluted peak at 6.2 minutes was present in all APRT2 assay fractions and could not be satisfactorily isolated and identified under current assay conditions.

## Conclusion

As a purine auxotroph, *Trypanosoma brucei ssp*. rely on purine phosphoribosyltransferases to salvage purines from their hosts for the synthesis of purine monophosphates. The APRT activity is important not only in adenine and adenosine metabolism, but also for its role in the activation of adenine-based pro-drugs [35, 37, 38]. All trypanosomes present two putative *aprt* genes in tandem; however, this characteristic is not observed on the other parasitic protozoa. APRT activity is unannotated or absent in *Plasmodium falciparum* [75], and *Toxoplasma gondii* [76]. Both *L. donovani* [77] and *G. lamblia* [78] present a single *aprt* gene, and both *Ld*APRT and *Gl*APRT share higher sequence conservation to *T. brucei brucei* APRT1 (52% and 35%) than to APRT2 (27% and 29%).

The trypanosomal APRT2 genes cluster apart from APRT1 on phylogenetic analyses and are more closely related to OPRTs than to other Type 1 PRTases [35]. However, we were unable to show enzymatic activity using orotate as substrate for the phosphoribosyl transfer, nor other purine bases known to be substrates of the Type 1 PRTases (S9 Fig and S10 Fig). We could not detect any enzymatic activity when APRT2 was tested as a possible adenosine phosphorylase, 5’-nucleotidase, or PRPP synthetase (S9 Fig and S10 Fig). The possibility of a PTM as a modulator of APRT activity was investigated, and APRT2 was shown to be unresponsive to MetO modifications as indicated by the lack of increased activity in the M to Q mutants (APRT2 M73Q, APRT2 M128Q, and APRT2 M156Q) (Fig 7 and S13 Fig). Some bacterial members of the Type 1 PRTases family seem to have no catalytic function, and instead act as expression regulators of genes involved in purine and pyrimidine synthesis [61]. Although a regulatory role for APRT2 cannot be ruled out at this moment, it is unlikely, as gene expression regulation in *Trypanosoma* is mainly achieved by post-transcriptional control [64], and the initiation of transcription by RNA polymerase II is not controlled for individual protein encoding genes [79].

The characterization of APRT1 activity here and in a recently published study [41], its detection in proteomics studies [24, 26], and higher sequence identity to other protozoan APRTs, allied to APRT2 lack of relevant catalytic APRT activity, indicate APRT1 is the only functional APRT enzyme in *T. b. brucei*. Although APRT1 is located in the cytosol [35, 41] while several other enzymes of the purine metabolism are glycosomal [23, 26]; the subcellular localization of the purine salvage enzymes is not conserved among the protozoans, nor is this pathway restricted to a single subcellular compartment [35, 80]. Our results reveal APRT1 has a greater preference for the forward reaction -in accordance with the parasite’s nature as purine scavengers. *T. brucei* purine transporters show great affinity for the free purine bases [31, 32], and the toxicity of high intracellular adenine concentrations is well described [35], highlighting the importance of phosphoribosyltransferases that greatly favor the forward reactions for incorporation of purine bases into nucleoside monophosphates. Our results provide a greater understanding of enzymes involved in adenine salvage, strengthening the ability to utilize APRT for therapeutic purposes in the development of drugs or pro-drugs of high selectivity and affinity. The role of APRT2, if any, in the purine salvage pathway, or as a Type 1 PRTase, requires further evaluation, while keeping in mind a possible event of gene duplication and loss of activity.

## Acknowledgements

The *P. pastoris* strain (smPP) and pPICZ vectors used to express APRT1 and APRT2 were a generous gift from Dr. Arthur Laganowsky. Samantha Schrecke provided assistance with establishing expression conditions for APRT1 and APRT2 in *P. pastoris*. Zahra Moghadamchargari collected the native mass spectrometry data for recombinant APRT2-Ntag and APRT2-Ctag.

